# Dynamic structural changes in wheat vegetative development as an adaptive response to drought stress

**DOI:** 10.1101/2025.08.07.669084

**Authors:** Agata Leszczuk, Nataliia Kutyrieva-Nowak

## Abstract

**Background and Aims:** The objective of our research was to define the precise structural response in wheat seedlings correlated with the duration of drought stress. For this purpose, we selected structural components of the cell recognised by specific molecular probes, which are molecules involved in rapid spatial cellular rearrangements: hydroxyproline-rich glycoproteins, xylan, and pectic compounds.

**Methods:** Using basic molecular techniques, we identified the transformations occurring within the cell and elucidated the mechanism triggered by growth in the absence of water.

**Results:** Our general observations are as follows: 1) remodelling of the cell wall after just 5 days of drought conditions; 2) organ-specific responses for drought resistance; 3) drought triggers the aggregation or cross-linking of molecules in the cell wall (appearance of larger molecular mass fractions) and causes degradation or breakdown of cell wall components (appearance of low molecular masses); 4) changes in the elemental economy due to modifications in cellular assembly.

**Conclusion:** Our finding of the deposition of un- and esterified homogalacturonans (HGs) and AGPs indicates reconstruction of cell wall as a means of prevention of drought effects. A stress-induced higher level of unesterified HGs permits calcium cross-linking, which enhances cell wall rigidity and helps in intracellular water preservation.

**Highlight statement:** Dynamic changes in wheat as a response to drought include remodelling of the cell wall after 5 days of drought,modification in the elemental composition, deposition of HGs, xylan, and AGP.

## INTRODUCTION

The inevitable intensification of extreme weather phenomena is the agenda of all international bodies for assessing the knowledge of climate change. According to data from NASA, Hadley, NOAA, Berkeley, Copernicus, and JRA-3Q, the Intergovernmental Panel on Climate Change (IPCC) and the World Meteorological Organisation (WMO) provided a synthesis of the global surface temperature records. It finds that 2024 was the warmest year on record and was the first in which average global temperatures at the planet’s surface exceeded 1.5°C above pre-industrial levels (Carbon Brief’s reports, 2025). A phenomenon undoubtedly associated with climate-related hazards and intensified by global warming is drought. Drought is among the most complex and fundamentally cross-disciplinary phenomenon with wide-ranging and cascading impacts across societies, ecosystems, and economies (Cook et al. 2018; Naumann et al. 2021).

Numerous research groups address the topic of drought and, according to the Web of Science database, over 15,000 scientific reports were published with the declared keyword ‘*drought*’ in 2024. What is clear is that extensive studies have shown the severe impact of drought on plants at the organ level (de Vries et al., 2020; Gupta et al., 2020; Valarmathi et al., 2023; Wang et al., 2024) and at the cellular level (Chen et al., 2020; Ezquer et al., 2020; Ganie and Ahammed, 2021). Conversely, plants have also evolved complex cell-type-specific processes to adapt to a dynamic environment (Cantó-Pastor et al., 2024). Studies suggest that plants exhibit distinct signals in response to emerging drought stress in both roots and leaves. To balance the limited amount of water, plants have developed numerous pathways, including increasing root water uptake from soil, decreasing water loss by leaves, and customising osmotic processes in tissues. The root system architecture undergoes morphological modifications to enhance abilities for water absorption, including longer roots with reduced branching angles and reinforced hydrotropism for optimisation of water acquisition, root hydraulic conductance increases, and changes in root exudation chemistry. In the case of leaves, alterations induced by drought include turgor-driven shape changes in guard cells, which are correlated with stomatal closure and rapid defence against dehydration as well as changes in cytoskeleton dynamics, remodelling of the cell wall, and plasma membrane stability (de Vries et al., 2020; Gupta et al., 2020). The cell wall-plasma membrane serves as a venue for sensing metabolic changes that affect plant development and stress responses (Vaahtera et al., 2019). The first symptom of the cell wall response to drought stress is controlled cell wall folding, which prevents tearing of the plasma membrane from the cell wall, which is essential to maintaining cell integrity (Chen et al., 2020). Maintaining cell wall integrity during dehydration involves two opposing effects: stiffening and loosening, which involve multiple components, from changes in the polysaccharide composition to differential RNA expression. This is due to the reorganisation of the plant cell wall and dynamic pectin changes, specifically higher levels of unesterified homogalacturonan upon drying. Next, a higher level of unesterified pectin effects an increase in the concentration of calcium in the cell wall, which induces large Ca^2+^ efflux from mesophyll cells, accompanied by K^+^ efflux and H^+^ influx (Chen et al., 2020). Furthermore, cell wall integrity is undoubtedly related to Hydroxyproline-Rich Proteins (HRGPs), including extensins and arabinogalactan proteins (AGPs) acting as ‘plasticisers’ maintaining the flexibility of cell walls in drought conditions (Chen et al., 2020). Also, the accumulation of these proteins involved in the resistance mechanism and the formation of a ‘buffer zone’ prevents direct interaction between membranes and the CW matrix (Ezquer et al., 2020).

In addition to cell wall structure and integrity, the presence of elements is crucial for maintaining cellular functions in water-deficit conditions. Water management in plants is linked to the presence and balance of essential elements that can enhance their tolerance to water stress. Some of them, known as ‘stress tolerance elements’, such as potassium (K), calcium (Ca), sodium (Na), and silicon (Si), provide a form of protection in drought conditions. Zhang and coworkers (2024) showed evidence that Na and Ca are the primary elements to combat drought stress through improvement of the activities of antioxidant enzymes (SOD, CAT, and POD) to alleviate oxidative stress. Na^+^ increases plant resistance to drought because it influences the regulation of the osmotic potential and cell turgor, accelerating stomatal closing during water deficiency (Zhang et al., 2024). Also, Ca is an essential structural component in plants that regulates cell wall rigidity via cross-linkage of pectin COOH—groups (unesterified) in the middle lamella (Kour et al., 2023). In drought conditions, it significantly increases antioxidant defence activity (Torun, 2022), maintains proper folding and translocation of proteins, and increases relative water content in leaves and cell membrane stability (Li et al., 2020). As a secondary messenger, Ca transmits drought signals, triggering a signalling cascade in guard cells that results in stomatal closure, thus reducing water loss. Along with the improvement in water status, there was a correlated minimisation of membrane damage by Ca, which appears to reduce the destructive effects of stress by increasing proline and glycine content (Song et al., 2008). Participation in the opening and closing of stomata, thus limiting the evaporation of water from plants, is also the role of K. This function is mainly due to its role in maintaining ion homeostasis, cellular integrity, cell membrane stability increase, and enzymatic activities (Hasanuzzaman et al., 2018). As a component of drought adaptation, K mediates xylem hydraulic conductance, cell turgor maintenance, stomatal movement, and sufficient gas exchange (Oddo et al., 2011). Another example is the mitigating effect of Si on drought stress that has been observed in many crop species, even in plant species that have a poor ability to accumulate Si (Wang et al., 2021). Si has been found to increase crop biomass in drought stress conditions, which is attributed to increased total root length, surface area, and plant height. At the cellular level, Si is known to increase root endodermal silicification and suberisation (Fleck et al., 2011); it also regulates the activities of enzymes involved in carbohydrate metabolism and affects the lignification of cell walls, regulating assimilate synthesis efficiency (Zhu et al., 2016). Additionally, Si creates mechanical barriers, increasing plant tolerance to soil water deficits by settling on plant cell walls and in intercellular spaces (Wang et al., 2021). Root endodermis silicification is associated with greater plant drought tolerance through Si deposition in the cytoplasm, leading to the formation of high molecular weight Si complexes in plant cell vacuoles (Malik et al., 2021).

Taking into account the large amount of data on changes occurring in the plant as a result of its growth in drought conditions, it was noticed that there is a lack of detailed and uniform characteristics of plant modifications, a step-by-step description of disorders appearing in the plant, with particular emphasis on the time of appearance of symptoms. Thus, the objective of our research was to define the precise structural response correlated with the duration of stress. For this purpose, we selected structural components of the cell recognised by specific molecular probes, which are considered molecules taking part in rapid spatial cellular rearrangements. We selected hydroxyproline-rich glycoproteins (HRGPs), arabinogalactan proteins (AGPs), xylan, and pectic compounds, which are key components involved in the organisation of the temporal-spatial assembly of each plant cell. Using basic molecular techniques, we marked the transformations occurring within the cell. Next, based on the collected data, we managed to develop a mechanism triggered by growth in the absence of water. Understanding the defence mechanisms of plants allows us to find solutions that enable the growth of plants under environmental stress. This is undoubtedly important in crop research, considering that the proposed protection solutions must be characterised by high precision and reliability.

## MATERIAL

Plants of winter wheat (*Triticum aestivum*) cv. ‘Arkadia’ were used as a research material. 5-day-old seedlings (stage named: start) were placed in soil for the next 5, 10, 15, and 20 days. Soil was collected from a wheat field (51°25′53.5” N, 19°37′05.3” E). During the experiment, one part of the plants was not watered (named: water stress – drought), and one part was treated as a control; thus, the plants were watered throughout the duration of the cultivation (Figure 1). All other growing conditions were the same in both cases. Every 5 days, materials – leaves and roots – were collected for research. For each sample variant, 50 plants were analysed and subjected to appropriate preparation. The materials were weighted, frozen in liquid nitrogen, and immediately homogenised for extraction and molecular techniques. In the case of microscopic analyses, the material was cut into 0.5 cm × 0.5 cm cubes and then processed using chemical fixatives.

**Figure 1.**
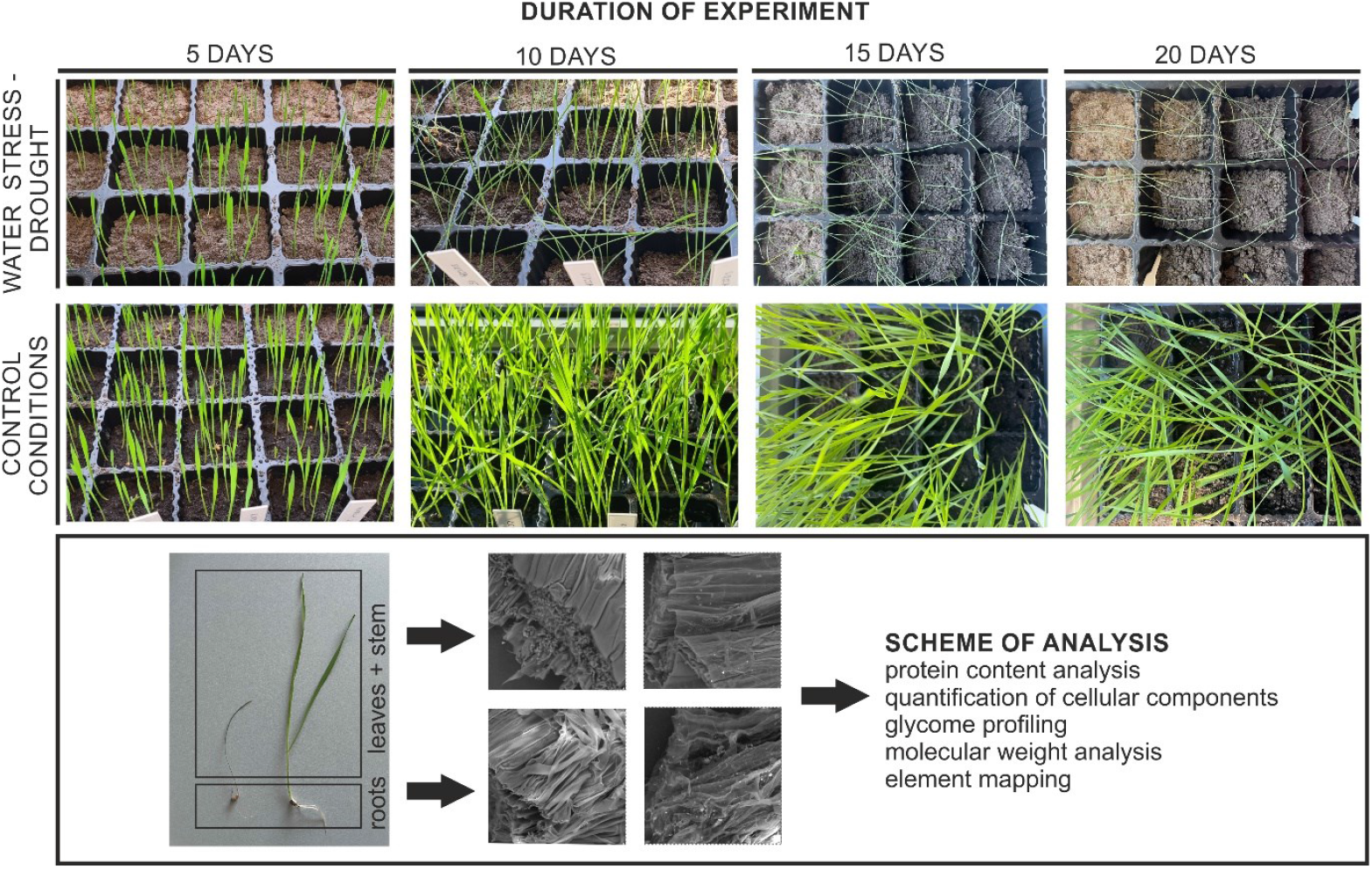
Experimental design ‘Plant cultivation in drought conditions’: soil drought pot experiment: after 5 days, seedlings were exposed to water stress conditions, and the following samples were collected every 5 days: 5, 10, 15, and 20 days of the experiment. A control experiment was conducted simultaneously with plants watered regularly. The research material consisted of leaves, stems, and roots of young wheat plants, which were then subjected to analyses: protein content analyses, quantification of cellular components, glycome profiling, molecular weight analysis, and elemental mapping.

## METHODS

Protein extraction and electrophoretic separation – Sodium Dodecyl Sulfate-Polyacrylamide Gel Electrophoresis (SDS-PAGE)

To determine the protein content in the samples, a standard protein extraction with Laemmli buffer (62.5 mM Tris-HCl (pH 6.8), 2% SDS, 2 mM EDTA, 1 mM PMSF, 700 mM β-mercaptoethanol, and protease inhibitors) was performed. First, the collected plant tissue was frozen in liquid nitrogen, homogenised with the buffer, and heated at 95°C for about 5 minutes to break disulphide bonds. Next, the samples were centrifuged (14,000 rpm), maintaining the rigor of 4°C for 20 minutes to ensure complete lysis. The supernatant was collected from the homogenate obtained, and its protein concentration was determined using the Bradford assay (Bradford, 1976).

The supernatant was then loaded into polyacrylamide gel for electrophoresis. The Sodium Dodecyl Sulfate-Polyacrylamide Gel Electrophoresis (SDS-PAGE) was carried out using a typical procedure involving several steps, including gel casting, running the gel, and protein visualisation with Coomassie Brilliant Blue staining. For the determination of molecular mass, Pierce™ Prestained Protein MW Marker was applied (Thermo Scientific, USA). The quantitative analysis of protein bands was performed using the GelDoc Go Imaging System (Bio-Rad, USA) with Image Lab Software version 6.1 (Bio-Rad, USA). Two distinct experiments were carried out.

### Immuno-dot blot assay

To quickly verify the content of individual components in the sample, detection was performed using Immuno-dot assays (IDAs). After the preparation of the nitrocellulose membrane, a 2 µL volume of each sample was pipetted onto the membrane in dots (in 3 replicates). The dots were dried, and then the typical immunocytochemical reaction procedure was started, including preincubation with BSA and incubation with the primary antibody at a concentration of 1:500, followed by a series of washes with TBST and incubation with the secondary antibody at a concentration of 1:1000. In our studies, 10 antibodies recognising specific epitopes that constitute the structural basis of each cell were used. Commercial antibodies from Kerafast (USA) were used. Their detailed specifications are described in the table below (Table 1). For visualisation, colorimetric detection was chosen, which allowed semi-quantitative interpretation of data based on colour depth analysis. Data analysis was performed using Image Lab Software v. 6.1 (Bio-Rad, USA) and then presented in the form of a heat map.

**Table 1.**
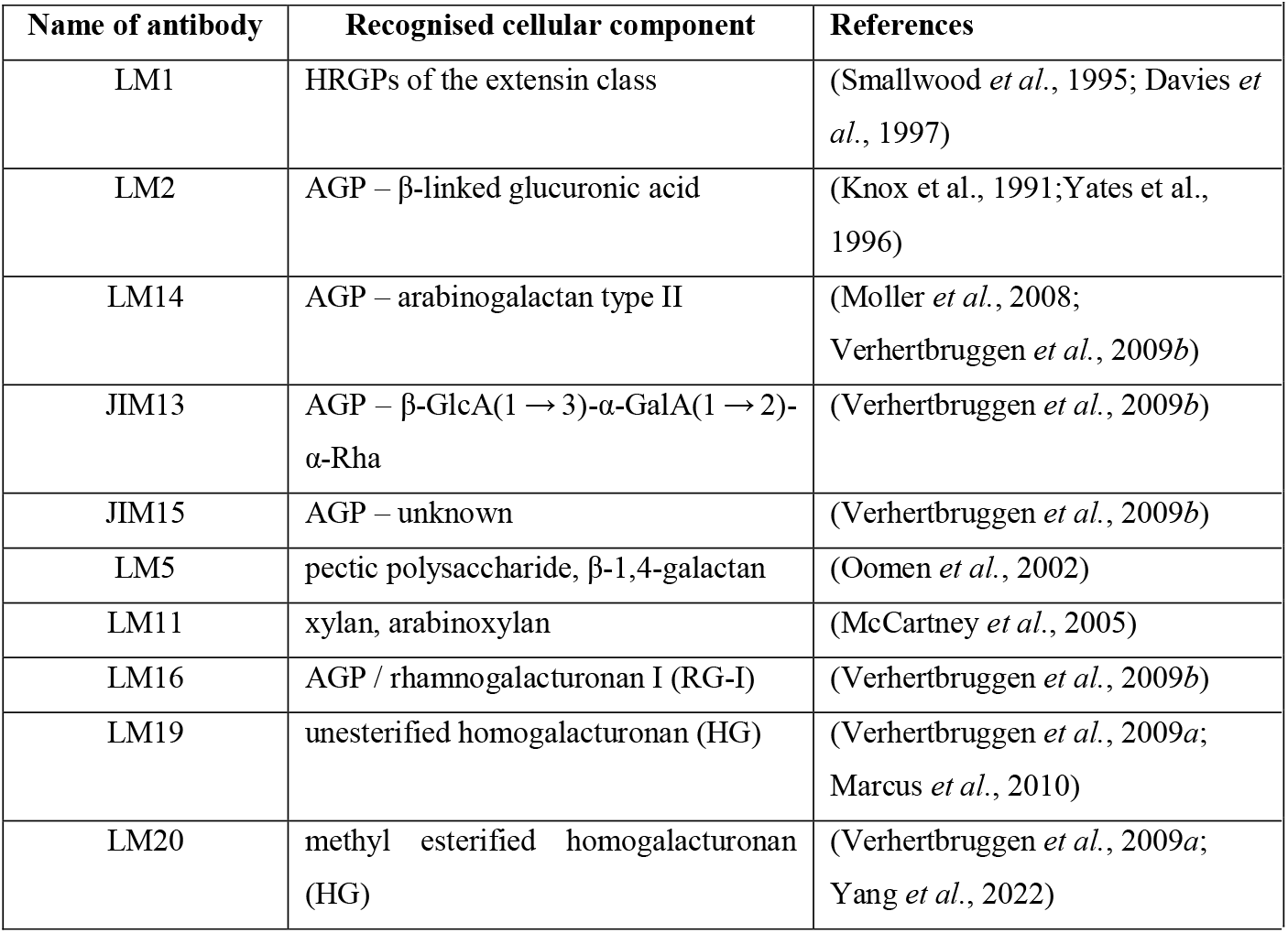
Antibodies used in the current research.

| Name of antibody | Recognised cellular component | References |
| --- | --- | --- |
| LM1 | HRGPs of the extensin class | (Smallwood *et al.*, 1995; Davies *et al.*, 1997) |
| LM2 | AGP – β-linked glucuronic acid | (Knox *et al.*, 1991; Yates *et al.*, 1996) |
| LM14 | AGP – arabinogalactan type II | (Moller *et al.*, 2008; Verhertbruggen *et al.*, 2009b) |
| JIM13 | AGP – β-GlcA(1 → 3)-α-GalA(1 → 2)-α-Rha | (Verhertbruggen *et al.*, 2009b) |
| JIM15 | AGP – unknown | (Verhertbruggen *et al.*, 2009b) |
| LM5 | pectic polysaccharide, β-1,4-galactan | (Oomen *et al.*, 2002) |
| LM11 | xylan, arabinoxylan | (McCartney *et al.*, 2005) |
| LM16 | AGP / rhamnogalacturonan I (RG-I) | (Verhertbruggen *et al.*, 2009b) |
| LM19 | unesterified homogalacturonan (HG) | (Verhertbruggen *et al.*, 2009a; Marcus *et al.*, 2010) |
| LM20 | methyl esterified homogalacturonan (HG) | (Verhertbruggen *et al.*, 2009a; Yang *et al.*, 2022) |

### Western immunoblotting assay

To characterise specific epitopes, a Western immunoblotting assay was carried out. The general step-by-step guide for Western blotting preparation is described in our previous paper (Kutyrieva-Nowak et al., 2023). Briefly, after separation of proteins using SDS-PAGE, Western Blotting involved wet transferring onto a nitrocellulose membrane (PVDF) (Thermo Scientific, USA). Immediately after blocking non-specific binding sites with BSA in PBS, the membrane was incubated with a primary antibody (Table 1), followed by a secondary antibody – Anti-Rat-IgG conjugated with alkaline phosphatase (AP) (Sigma-Aldrich, USA). The bands were visualised using colorimetric substrates – 5-bromo-4-chloro-3-indolyl phosphate (BCiP; Sigma, USA), nitro-blue tetrazolium (NBT; Sigma, USA), and N, N-dimethylformamide (DMF; Thermo Scientific, USA). The band thickness and colour depth were used for qualitative analysis at three ranges of molecular weights – high (120-60 kDa), medium (60-25 kDa), and low (25-20 kDa) molecular ranges. The quantitative analysis of protein bands was performed using Image Lab Software v. 6.1 (Bio-Rad, USA).

### Glycome profiling

For both the qualitative and quantitative identification of structural components, ELISA on a 96-well plate (Nunc MaxiSorpTM flat-bottom, Thermo Fisher Scientific, Denmark) was carried out. After immobilisation at 37°C for 72 h, the coated plate was washed with PBS (pH 7.4), preincubated with 0.1% BSA, and incubated with primary antibodies (Table 1) at a concentration of 1:20 and Anti-Rat-IgG alkaline phosphatase (AP)-conjugated secondary antibody in a dilution of 1:500. The reaction was developed with a solution of p-nitrophenyl phosphate (PNPP) and stopped with 2M NaOH. The absorbance was measured in an ELISA reader (MPP-96 Photometer, Biosan) at 405 nm. The AP activity was measured by the release of PNP from PNPP. The concentration of PNP was determined using a calibration curve (y = 0.0808x, R^2^ = 0.9854). Three repetitions were conducted. Analysis of variance (one-way ANOVA) and Tukey’s Honestly Significant Difference (HSD) post hoc test (Statistica v.13 tools (TIBCO Software Inc. USA)) were used to compare the mean results. For all analyses, the significance level was estimated at p<0.05.

### Scanning Electron Microscopy with Energy Dispersive Spectroscopy (SEM-EDS)

In order to suppress any changes occurring in the tissue during morphological observations, cut leaf and root fragments were subjected to chemical fixation. Initially, samples in 2% paraformaldehyde with 2.5% glutaraldehyde were de-aired in a vacuum pump and then dehydrated in a series of alcohol (10-99.8% ethanol) and acetone. Such fragments were placed on the SEM stub with carbon tape. The morphology of tissue was observed using a scanning electron microscope (SEM, Phenom ProX, Pik Instruments). Images were taken at the same magnification, around 2.000 kx, at an accelerating voltage of 15 kV with a BSD detector. In turn, for elemental analysis, the applied SEM microscope was equipped with an energy dispersion X-ray spectrometer (EDS). Some samples appeared as areas with more intense colours in the elemental maps, showing a higher concentration of specific elements (Na, K, Si, Ca). The data obtained from three replicates were used for a comparative analysis of the content of selected elements.

## RESULTS

The analyses were carried out on research material collected over 20 days. Clearly, during cultivation with no access to water, the plants were different after 10 days from the control plants (watered). Each subsequent material, on the 15th and 20th day of drought, was increasingly dried out (Figure 1). The morphological analysis of the plants did not reveal any difference in the material cultivated on day 5 (Figure 1), but the molecular and biochemical analyses indicated the first signs of response of the plant leaves and roots to the drought stress.

The protein content in the leaves of 5-day-old seedlings was 250 µg/mL, and then, during plant growth in control conditions, it increased to about 300 µg/mL and remained at the same level for 20 days. In the drought conditions, the protein content decreased after just 5 days, reaching a value approximately six-fold lower than in the leaves of the control plants. This low protein level in the leaves decreased slightly as the experiment progressed, and after 20 days, the protein content was below 50 µg/mL. In the case of roots, the protein content was slightly lower than in the leaves; it was around 80 µg/mL in a 5-day-old seedling and almost 250 µg/mL after 5 days of growth, and then it remained below 200 µg/mL throughout the growth of the plant. The drought conditions affected the protein content in the root; its value decreased below 100 µg/mL after only 5 days of cultivation and was maintained at the same level during the next days of cultivation (Figure 2).

**Figure 2.**
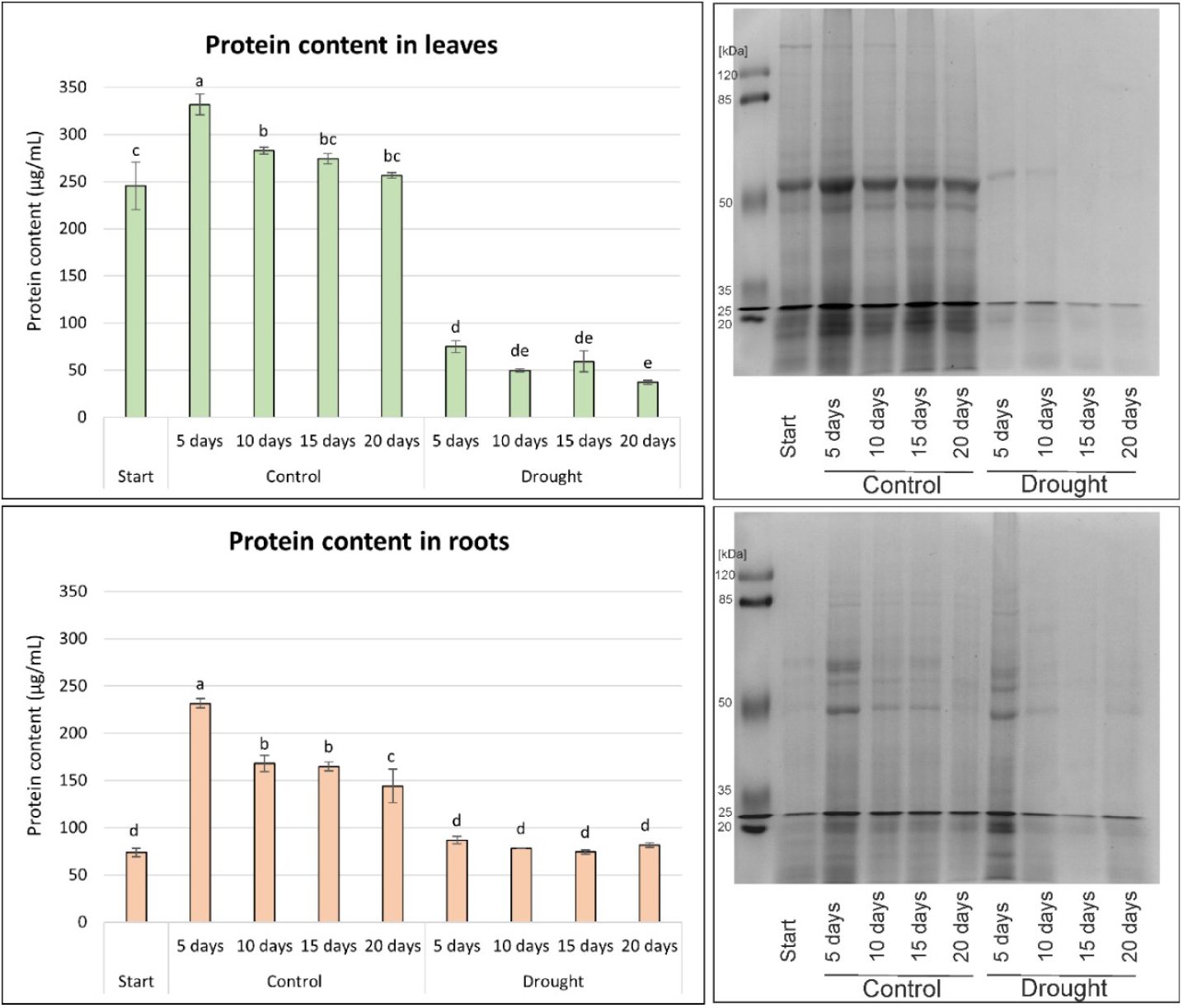
Quantification of protein content in wheat seedlings. Bradford method and analysis of their molecular weights by electrophoretic separation and Coomassie blue staining. The concentration [µg/mL] was calculated by mean absorbance values (n=3 biological replicates for each sample) compared to a standard curve created using BSA. Statistical significance was determined using a Tukey post-hoc test (different letters indicate a significant difference at p < 0.05).

After performing gel electrophoresis, several distinct bands were observed in the leaf sample. The bands were most clearly visible, indicating a high abundance of proteins with a molecular mass of about 50 kDa. Much smaller bands with a molecular mass of 45 kDa and in the range of 20-30 kDa were also noted. Such a separation of proteins was characteristic of leaves of plants grown in control conditions. The visualisation of the electrophoretic separation of proteins isolated from the leaves of plants growing under drought was completely different. The bands indicating the presence of protein with a mass of about 50 kDa were visible only in the sample from leaves from the 5th and 10th day of the experiment (Figure 2). Separation of proteins isolated from roots allowed identification of molecules with molecular masses of about 60 kDa, 50 kDa, and 20 kDa (Figure 2). Moreover, the highest intensity of the bands was observed in roots after 5 days of growth. Proteins isolated from roots of plants growing in the drought conditions were characterised by molecular mass in a similar range. Also, the conditions of 10, 15, and 20 days of drought affected their presence and molecular mass, both of which decreased, indicating degradation of the molecules (Figure 2).

The immuno-dot blot method was employed for semi-quantitative analyses of structural alterations in plant organs as a response to the drought stress (Figure 3). Analysis of changes in the content of HRGPs, AGPs, pectins, and arabinoxylans was performed using the appropriate antibodies. The colorimetric intensity of dots is the first visual cue that indicates the relative amount of target molecules in an examined sample of leaves and roots. Higher colour intensity reflects a higher target concentration, while weaker or no colour signals indicate lower or absent target levels. The analysis showed that AGP (JIM13, JIM15, LM2, LM14) and other HRGPs, represented by extensins (LM1), remained at the same level in both leaves and roots during growth in the drought conditions. In contrast, a significant increase in the pectin (LM5, LM16, LM19, LM20) and xylan (LM11) concentrations was observed in both examined organs (Figure 3). A detailed analysis of quantitative changes in the plant requires a more detailed approach; therefore, an ELISA analysis was performed (Figure 4).

**Figure 3.**
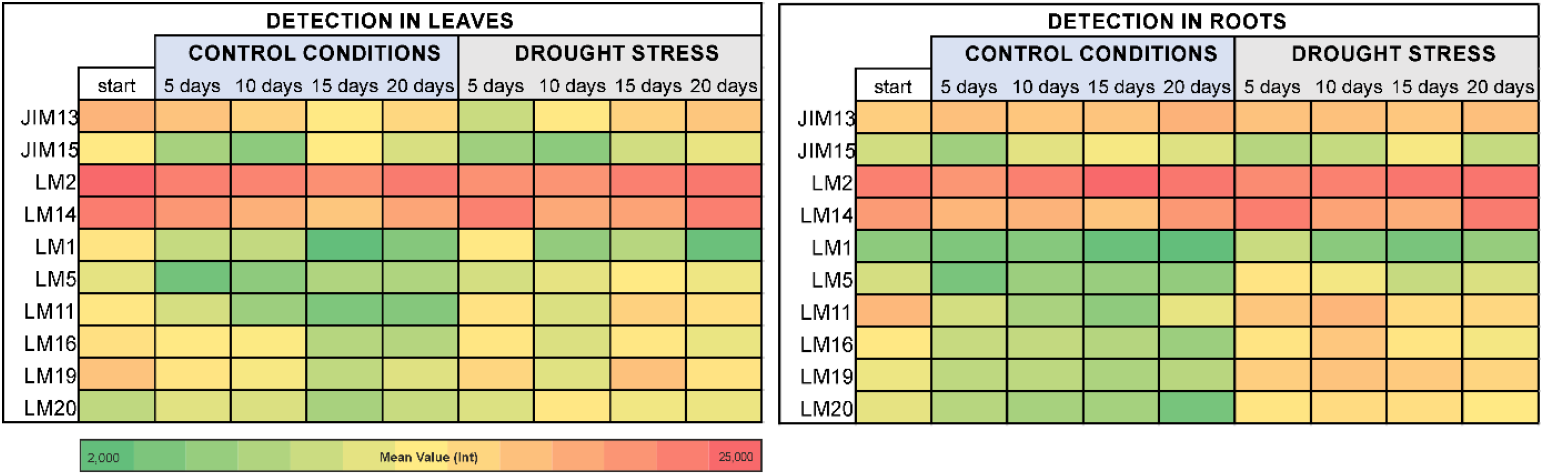
Semi-quantitative determination – immune-dot analysis of individual structural components in the analysed material: leaves and roots of wheat seedlings. Comparative analyses between the beginning (start) of the experiment and the cultivation for 5, 10, 15, and 20 days in the control conditions and under the drought stress. Dot blotting with JIM13, JIM15, LM2, LM14, LM1, LM11, LM5, LM16, LM19, and LM20 antibodies. Mean spot value of labelling intensity (n=3 biological replicates). Data represent the units of intensity (colour depth) – an indicator of the amount of target molecules in the sample.

**Figure 4.**
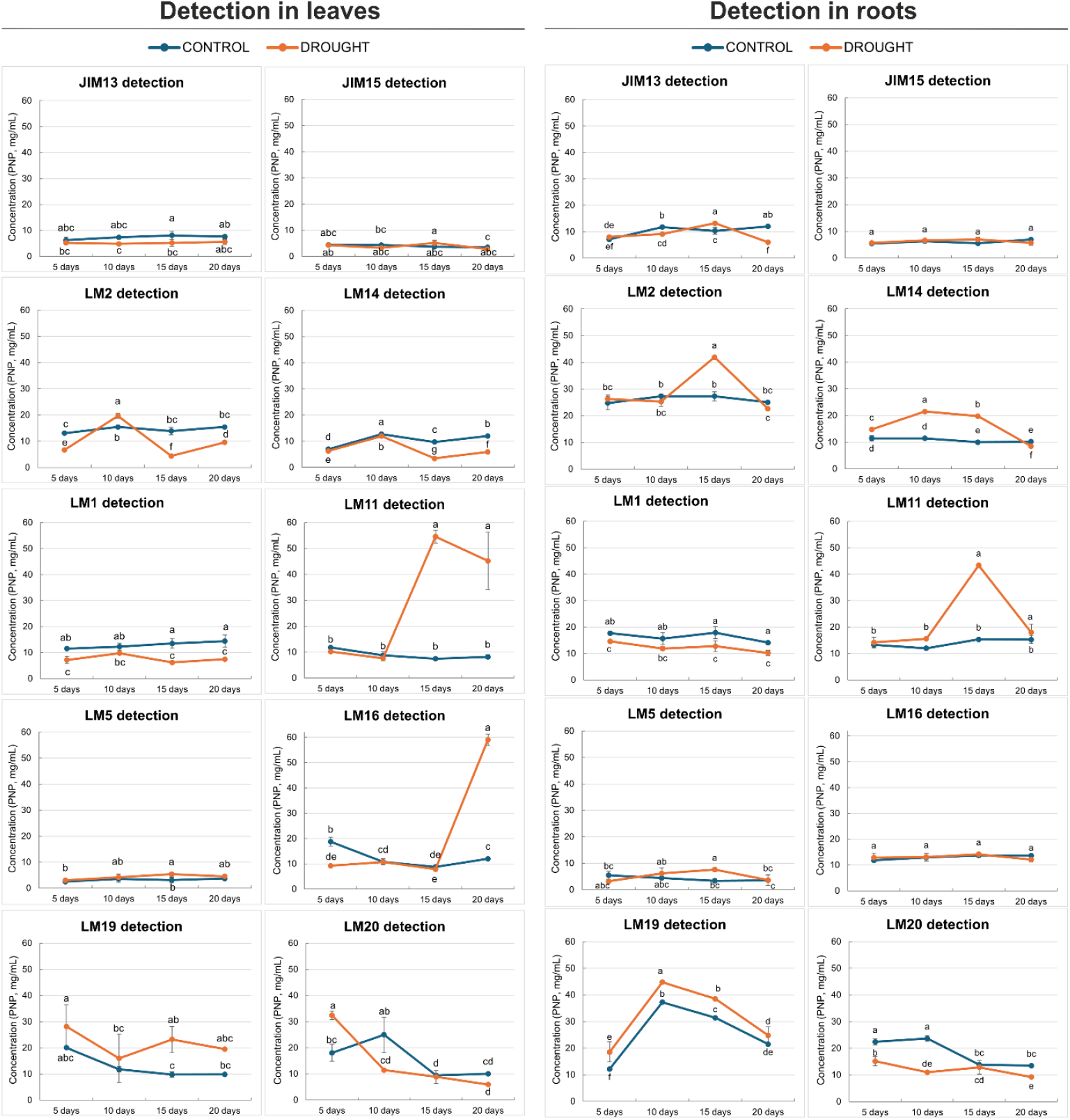
Quantification of individual structural components in the analysed leaves and roots of wheat seedlings. Comparative analyses between the cultivation for 5, 10, 15, and 20 days in the control conditions and under the drought stress. ELISA with JIM13, JIM15, LM2, LM14, LM1, LM11, LM5, LM16, LM19, and LM20 antibodies. The concentration [mg/mL] was calculated by mean absorbance values (n=3 biological replicates for each sample) compared to a standard curve created using PNP (p-nitrophenyl phosphate). Statistical significance was determined using a statistical Tukey post-hoc test (different letters indicate a significant difference at p < 0.05).

An ELISA test was performed to investigate the quantitative relationships between the content of various structural components and the occurrence of drought stress over the studied period (Figure 4). The components constituting the polysaccharide base of each cell were analysed and were identified as specific epitopes bound by antibodies, specifically HRGP – LM1; β-linked glucuronic acid – JIM13, JIM15, LM2, LM14; arabinoxylan – LM11, β-1,4-galactan – LM5; RG-I – LM16, unesterified HG – LM19, and esterified HG – LM20. The amount of individual compounds was calculated as the concentration of PNP, resulting from the breakdown of PNPP, and expressed in [mg/mL]. Quantitative changes, caused by water deficiency during growth, were observed in the leaves and roots.

The detection of the aforementioned components in the leaves revealed differences between the control plant and the drought stress variant, with the most pronounced changes observed in the case of LM2, LM1, LM11, LM16, LM19, and LM20 antibodies. In the case of the AGP analysis, trace changes were observed in the reactions with JIM13 and JIM15. In both cases, the concentrations did not exceed 10 mg/mL. However, the reaction with LM2 indicated an increase in the AGP content on the 10th day of drought (20 mg/mL), followed by a sharp decline below 10 mg/mL, which persisted throughout the subsequent stages of the experiment. A similar trend with a decrease in AGP levels was noted using the LM14 antibody. The analysis of extensins showed a decrease in their content due to the drought stress, starting from the beginning of the experiment on day 5 and continuing until day 20 of the drought. The most pronounced response to the stress conditions was observed in the concentration of the LM11 epitope, i.e., arabinoxylan. While the concentration after 10 days of the experiment remained at the same level as in the control leaves (around 10 mg/mL), on day 15 of the experiment, the concentration increased more than fivefold (55 mg/mL) and remained elevated throughout the subsequent stages of the experiment (45 mg/mL). The quantitative changes in pectin concentrations were not uniform for all the analysed types. No changes were observed in LM5-recognised β-1,4-galactan (not exceeding 10 mg/mL). However, the RG-I concentration on day 20 of drought increased more than fivefold (55 mg/mL), compared to the control leaf RG-I concentration (12 mg/mL). The native concentration of HGs was also disrupted due to water deficiency. Interestingly, depending on the level of esterification, their concentration changed in different ways. The concentration of unesterified HG increased from the early days of the experiment (18-28 mg/mL), compared to the control (9-20 mg/mL), while the concentration of esterified HG decreased from 32 mg/mL to 5 mg/mL starting from day 10 of drought (Figure 4).

Interestingly, changes were also observed in the roots, but they were quite different. The concentrations of AGP recognised by JIM13 and JIM15, β-1,4-galactan – LM5, and RG-I – LM16 were very similar in both roots grown under drought and in the control conditions. In the case of the roots, a different plant response was observed related to AGP recognised by LM2, as the concentration in roots grown for 15 days under drought increased to 40 mg/mL, compared to the control (below 30 mg/mL). Similarly, the concentration of AGP recognised by LM14 increased by more than twofold during the whole time of the experiment. The content of extensins recognised by LM1 in the roots, similar to the leaves, decreased due to the drought conditions. Interestingly, also in the root tissue, the concentration of arabinoxylan recognised by LM11 on day 15 of drought increased more than threefold (from 15 mg/mL to 43 mg/mL), showing a very clear peak as a potential response of the plant to the ongoing stress. Another similarity is the response of pectin compounds, whose native concentrations were also disrupted due to drought. Similarly, the concentration of unesterified pectins (LM19) increased, while esterified HG (LM20) decreased in comparison to the control roots. The ELISA results indicated that the experimental conditions caused significant alterations in the levels of the target molecules, compared to the control plants, suggesting a structural response to the drought (Figure 4).

In order to identify the cause of the differences in the concentrations of individual components, electrophoretic separations of the protein were performed to record the molecular mass values and their potential changes (Figure 5). It should be emphasised that the electrophoretic separation of the sample and subsequent Western blotting detection were performed on components that could potentially bind to proteins, i.e. AGP. Initially, variations in molecular mass and the emergence of single bands were noted, indicating that drought influences the formation or breakdown of the analysed molecules. Additionally, the band intensity varied in the different conditions examined. Figure 5 shows a summary of all membranes, with the most significant changes highlighted by a red frame.

**Figure 5.**
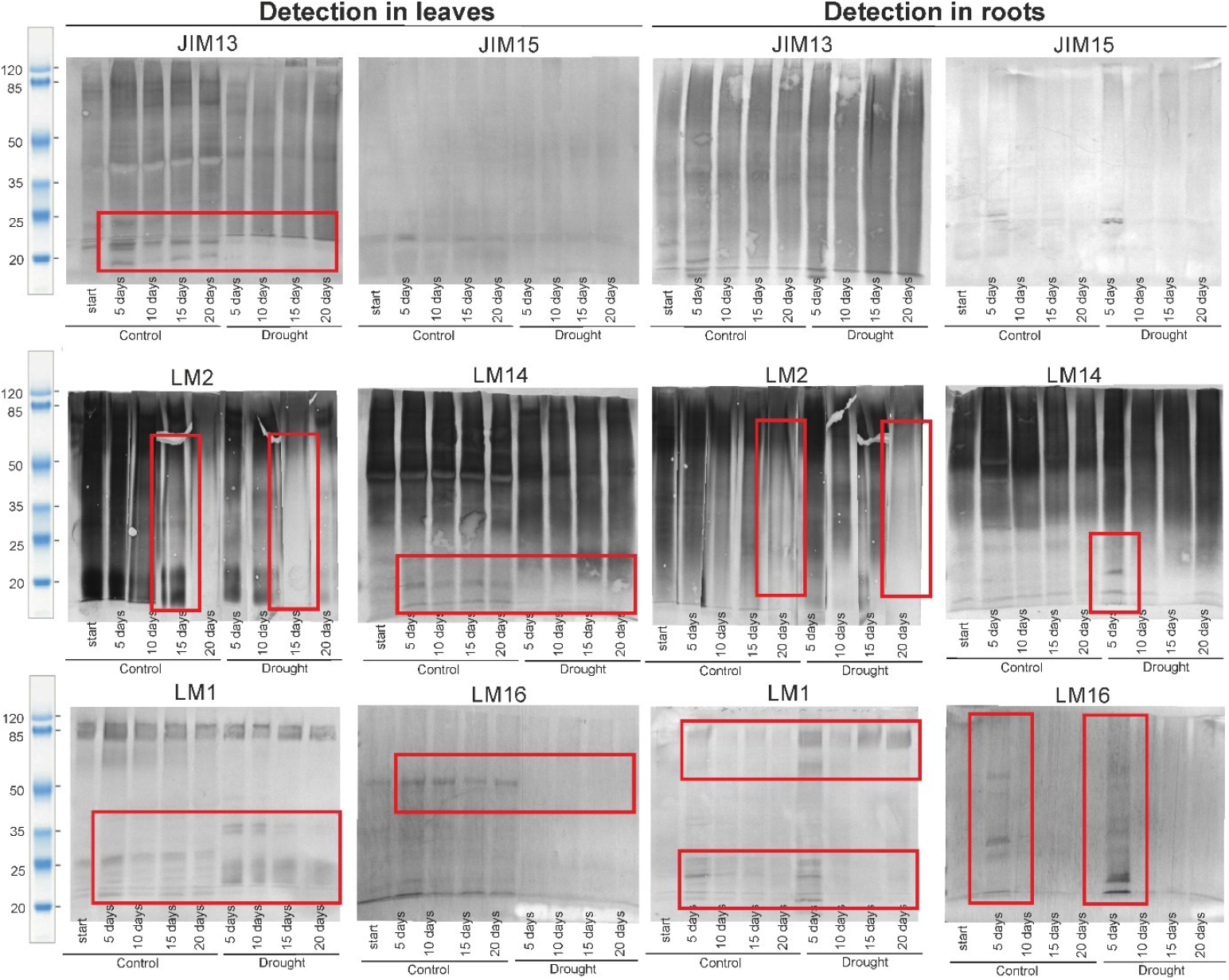
Electrophoretic separation with detection of individual structural components of the analysed leaves and roots of wheat seedlings. Comparative analyses between the beginning (start) of the experiment and the cultivation for 5, 10, 15, and 20 days in the control conditions and under the drought stress. Western blotting with JIM13, JIM15, LM2, LM14, LM1, and LM16 antibodies.

Interestingly, changes were observed in both leaf and root samples, but they were quite different. The changes observed in the leaf sample seemed to persist throughout the drought experiment, *i.e*., a loss of a single fraction was noted on day 5, and this change persisted until day 20 of the experiment. In the case of the JIM13 antibody, a loss of molecular masses below 20 kDa was observed from the onset of the drought conditions. Interestingly, the LM2-labeled AGP was not identified after 15 days of drought, and the sample only contained molecules with molecular masses above 60 kDa. Extensins also changed their molecular properties due to the drought – in the control sample, epitopes with a mass of around 25 kDa were present, but these were not identified in the drought-stressed plant samples. Additionally, in the sample from the leaves of non-watered plants, significant amounts of LM1 fractions below 25 kDa were observed as well as single bands on days 5 and 10 of drought in the range of approximately 35 kDa.

In the case of the root tissue analysis, changes in the molecular properties of AGPs were also observed. After 20 days of drought, no fractions below 85 kDa were present for LM2. However, for LM14, a clear increase in the amount of AGPs with lower molecular mass, i.e., below 35 kDa, was evident at every drought stage. Very interesting results were obtained for extensins, as the drought induced the appearance of bands indicating the presence of epitopes with a mass of 85-100 kDa. Furthermore, 5 days of drought triggered the synthesis of extensins, which were identified as single, very distinct bands in the range of 20-30 kDa. Intriguing results, providing new insights into the obtained data, were achieved through the AGP/RG-I analyses, specifically LM16. The root samples, already on day 5 of drought, exhibited a completely different molecular distribution of fractions compared to the watered plant samples. Bands at 35 kDa were identified, which served as a distinct marker not observed in the control samples.

The fraction distribution in each sample was quantified by measuring the relative intensity of the protein bands using Image Lab Software v. 6.1 (Bio-Rad, USA). The results obtained were presented as the fractions of components that constitute a single sample, enabling the evaluation of degradation or the emergence of new molecular fractions (Figure 6). The proposed analysis method highlights the effect of drought conditions on the decomposition of the cell wall. It is worth noting that a different distribution and its changes due to drought were demonstrated in the leaves and roots. Nevertheless, it can be unequivocally stated that drought has a significant influence on the molecular properties of the analysed components. In most of the analysed cell wall components, a reduction in fractions of compounds with molecular masses between 60-120 kDa was induced by the drought conditions. This applies to all selected epitopes. In the case of JIM15 and LM16 complete disappearance of the aforementioned fraction was noted, along with a simultaneous increase in compounds with lower molecular mass. A decrease in high molecular weight fractions was also observed with longer exposure to the drought stress, for example, in the reactions with JIM13, JIM15 and LM1.

**Figure 6.**
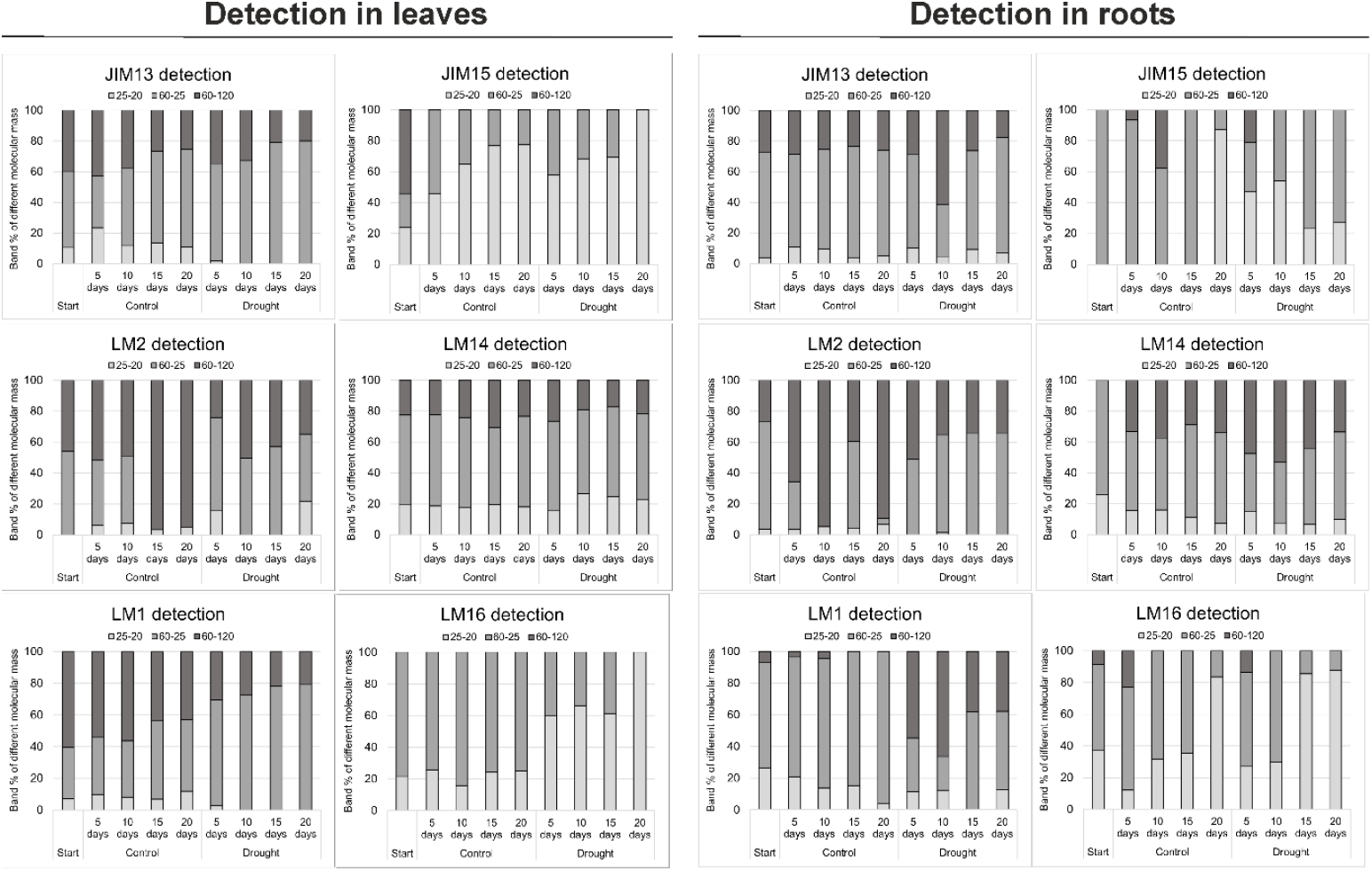
Determination of molecular mass values of individual structural components and their changes in the analysed leaves and roots of wheat seedlings. Comparative analyses between the beginning (start) of the experiment and the cultivation for 5, 10, 15, and 20 days in the control conditions and under the drought stress. Western blotting with JIM13, JIM15, LM2, LM14, LM1, and LM16 antibodies was based on the analysis of bands % in three molecular mass ranges: 20-25 kDa, 25-60 kDa, and 60-120 kDa. Results presented as % of particular molecular mass constituting the sample.

The EDS spectrum of the samples confirmed the presence of the primary elements in the examined tissues, including Na, Si, Ca, and K (Figure 7). The elemental composition data confirmed that plants growing in the controlled conditions, regularly watered, did not exhibit significant changes in the composition of both leaves and roots during the 20 days. These findings also provide valuable insights into the different timing of reactions or adaptations required for both roots and leaves in drought. Si predominated, while the other three analysed elements constituted a much smaller portion. After just 5 days of drought, there was a complete rearrangement of the elemental maps in the leaves. A significant decrease in Si was noted, accompanied by an increase in Na. After 20 days, a clear increase in the sodium content was identified. In the leaf analysis, no disturbances were observed in the level of K induced by the drought stress. In the case of the root tissue, the control sample also showed a predominance of Si and Na. After 5 days of drought, the Si content in the root decreased in the sample, while the levels of K and Ca increased. After 20 days of drought, the highest level in the root was recorded for Na, while the Ca content dropped to the level observed in the control sample.

**Figure 7.**
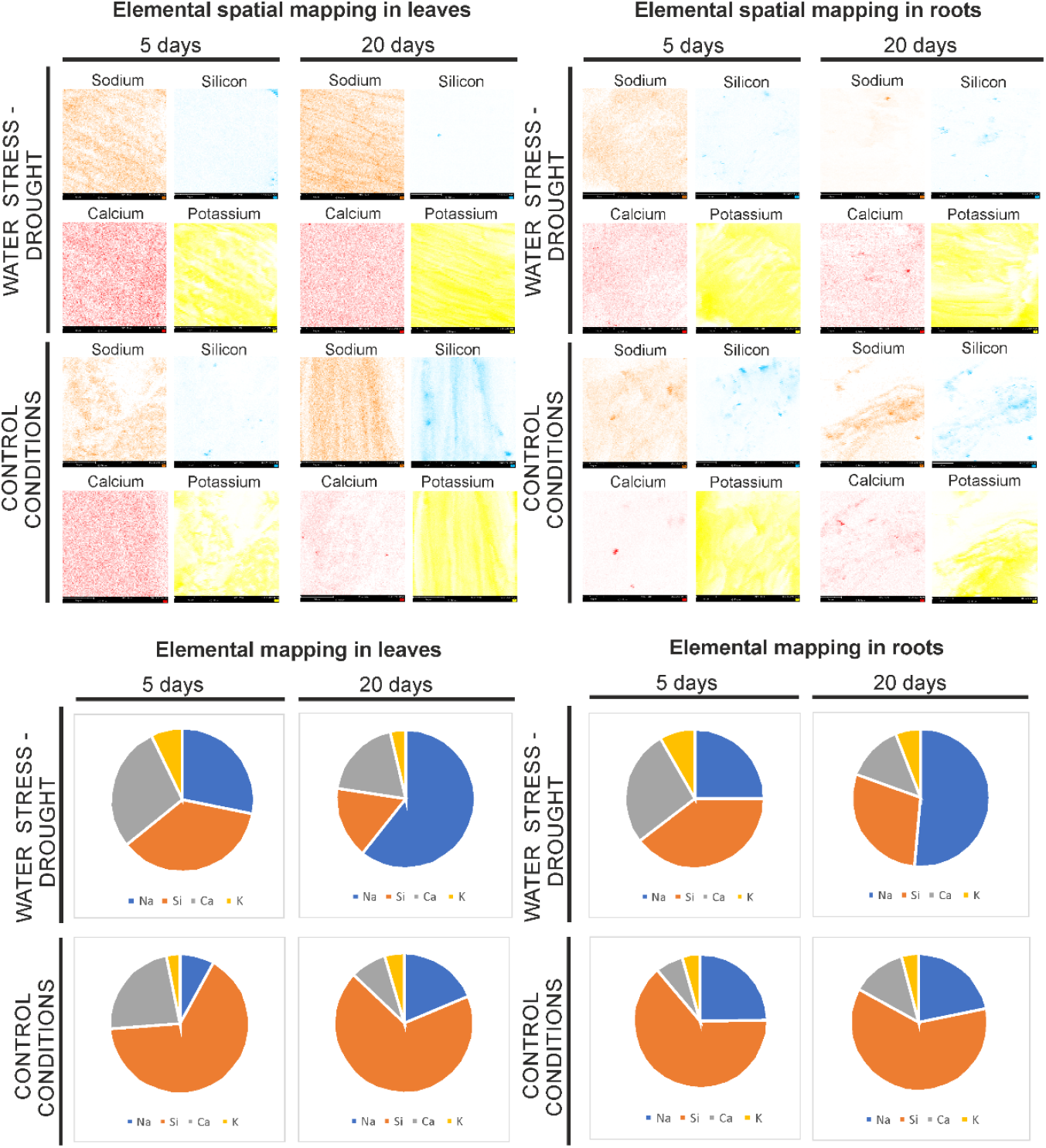
Spatial distribution of Na, Si, Ca, and K in the leaves and roots using elemental mapping with SEM-EDS. Comparative analyses between the cultivation for 5 and 20 days in the control conditions and under the drought stress. Qualitative analysis indicating regions rich in Na, Si, Ca, or K visualised as areas with higher colour intensity. The quantitative analysis was performed using SEM-EDS by analysing the X-ray peak intensities.

## DISCUSSION

Plants have developed adaptive mechanisms that enable them to respond effectively to stress factors, including drought. Drought stress is initially sensed by plant roots, which subsequently triggers the transmission of stress signals to the above-ground parts. Also, research shows that each organ and individual cells react to drought stress in unique ways (Ganie and Ahammed, 2021; Kim et al., 2024). Furthermore, these responses are closely correlated with the duration of such stress conditions. The growth conditions of plants, especially cereals, have an enormous influence on the quality of the crop. Our study aimed to gather data on the structural changes occurring in wheat plant cells as a result of drought to systematically determine the mechanism that governs the protection of both leaves and roots under environmental stress caused by drought conditions.

In the literature, there is ample information about plant adaptations to unfavourable conditions with water deficit (Moore et al., 2008; Tenhaken, 2015; Valarmathi et al., 2023). These are often molecular studies on model organisms with a strong emphasis on proposing genetic modifications that will allow them to overcome water-deficit stress. The following changes in the root are often distinguished: upregulation of xylan, arabinogalactan, and expansin in cell walls; upregulation of secondary metabolites involved in the biosynthesis of phenylpropanoids, carotenoids, flavones, anthocyanidins, and lignin; upregulation of genes involved in the transport of calcium, iron, and potassium; upregulation of protein kinases; abundance of abscisic acid (ABA) and auxin (Valarmathi et al., 2023). The molecular signalling cascade in the root in response to drought includes changes associated with cell wall loosening (Ganie and Ahammed, 2021). This, in turn is associated with the modification of the expression of genes related to the metabolism of cell wall proteins (GRP and UDP-GlcNAc pyrophosphorylase) and polysaccharides (OsCslF2, GMPase, xylose isomerase, and beta-1,3-glucanase). In the studies reported by Yang and coworkers (2006), overexpression of genes was determined as a response to drought stress; these genes are related to the metabolism of polysaccharides encoding enzymes that catalyse the synthesis of hemicellulose and β-glucan and the activation of mannosyl units (Yang et al., 2006). Also, genes encoding AGPs are reported to be involved in the drought-induced altered root phenotype, i.e., cell expansion and cell wall plasticization-mediated root growth (Yang et al., 2006).

In turn, in *Arabidopsis* leaves, drought stress has a strong negative effect on cell division, which is associated with reduced amino acid content (Skirycz and Inzé, 2010). On the other hand, in leaf tissues of three different wheat cultivars exposed to drought stress, increased expression of genes involved in amino acid biosynthesis was shown (Bowne et al., 2012). Thus, protein content is well correlated with species specificity. Furthermore, it is proven that glycophosphatidylinositol (GPI)-anchored membrane proteins are involved in drought tolerance via rice leaf rolling by regulating cell wall formation and leaf epidermal integrity (Li et al., 2017). The effective response of plants to drought also depends on the efficient upregulation of carbohydrate metabolism (Stitt et al., 2007). The expression of various genes coding carbohydrates synthesis, involved in cell wall formation and expansion, is increased by drought stress, i.e., the pectinesterase inhibitor gene, the 1,3-β-glucan synthase gene, and the xyloglucan galactosyltransferase gene (Verelst et al., 2013). Furthermore, shoot and leaf growth under drought stress is a result of a higher sucrose concentration, which acts as a signal for the upregulation of genes of OsCIN cell wall invertase, which metabolises sucrose to hexoses to maintain sink activity (Luquet et al., 2008). Interestingly, another conclusion is that reduction in cell wall pectin content, which affected cell wall integrity and the transpiration rate, resulted in increased drought sensitivity (Liu et al., 2014).

The object of our research is wheat, a cereal of great economic importance. Unfortunately, there are much fewer molecular studies on this research material, which is associated with the need to fill this gap in knowledge. Our research encompassed analyses at the cellular and molecular levels, taking into account all detailed structural changes in the cell wall architecture of wheat seedlings. Our general and collective observations are as follows: 1) remodelling of the cell wall after just 5 days of drought conditions; 2) organ-specific responses for drought resistance; 3) drought triggers the aggregation or cross-linking of molecules in the cell wall (appearance of larger molecular mass fractions) and causes degradation or breakdown of cell wall components (appearance of smaller molecular masses); 4) changes in the elemental economy due to modifications in cellular assembly. In addition to general information, our data provide specific data on changes in protein content. Namely, a clear decrease in the protein content was noted in the leaves after only 5 days of growth under the drought stress, and then a decreasing trend was still noted, compared to the control. In the case of the root, the protein content decreased similarly, but the amount remained at the same level over the subsequent days of the experiment. The glycome analysis performed here on both leaves and roots using an ELISA-based method revealed significant alterations in the levels of specific polysaccharides, which is consistent with the changes in molecular mass perturbing specific molecules. The immunocytochemical analysis of the leaf tissue showed that the epitopes of HGs were more abundant in the ‘dried’ tissue within just 5 days of the experiment, whereas no changes were observed in the epitopes of HRGPs, such as those of AGPs, β-linked glucuronic acid, and β-1,4-galactan. Interestingly, the concentration of unesterified HGs remained at a consistently higher level than the control for 20 days, whereas highly-esterified HGs decreased. Thus, this supports the well-documented idea that reduced methylation of pectins may result in a higher water holding capacity of the cell wall (Willats et al., 2001). Moreover, the amount of RG-I and arabinoxylan increased more than fivefold only after the longer exposure to stress, i.e., after 15 and 20 days of drought. In the root, a very similar trend of changes was noted in the case of pectic compounds, but the content of AGP recognised by LM2 and LM14 antibodies was different. In this case, significantly higher concentrations of AGP were determined in comparison to the root of plants developing in the control conditions. These disorganisations in the cell wall are in agreement with the major findings of studies in three wheat cultivars carried out by Rakszegi and coworkers (2014). In the study, in plants exposed to stress conditions, an increase in the side chains of pectic polymers RG-I and arabinoxylan was noted (Rakszegi et al., 2014). Our finding of the deposition of un- and highly esterified HGs and AGP may mean that the reconstruction of the cell wall to prevent the effects of drought is in agreement with studies on *Brassica napus* roots (Saja-Garbarz et al. 2024). The observed changes were connected with a manifestation of tissue strengthening in drought-stress conditions, which could counteract changes in turgor (Saja-Garbarz et al. 2024). Moreover, in studies on maize, the increase in arabinan-rich polymers, including AGPs, was explained as a contribution to more efficient water use during drought (Calderone et al., 2024). Immunodetection assays carried out by Calderone and coworkers (2024) showed an increase in unesterified HG upon drought, which was justified by increasing methylation of HG as indicative of either de novo deposition of newly synthesised HG or a decrease in the activity of pectin methyl-esterases (PME) (Calderone et al., 2024; Zhao et al., 2024).

Cell wall tightening and solute accumulation appear to be common phenomena in response to drought. In addition, drought stress affects non-covalent intra- and interpolymer bonds in the cell wall and alters the balance between wall extensibility and turgor pressure (Moore et al., 2008). This information is in agreement with the results obtained in our wheat studies. To determine the impact of drought on the cellular composition with emphasis on changes in the molecular mass of individual fractions, we performed Western blotting with detailed data analysis. The appearance of larger molecular mass fractions in the drought conditions suggests that drought triggers the aggregation or cross-linking of certain molecules in the cell wall. Altogether, these data suggest a stress response, where the plant attempts to reinforce its structure by modifying the molecular composition of its cell wall. In turn, the appearance of a lower molecular weight due to drought may indicate the disintegration of some cell wall components. This suggests that, in response to drought stress, the plant breaks down some of its structures, possibly to adapt to the conditions of limited water availability by releasing smaller molecular fragments that may be more mobile. In the future, more detailed studies will be needed to clarify the impact of loss of water on specific polysaccharide structure/interactions and the general impact on the cell wall organisation in root and leaf cells.

Another report in the context of changes in drought as a potential adaptive change is the modification in the elemental composition. The so-called ‘stress-tolerant elements’ stimulate enzymatic activity in the roots of plants grown in drought conditions, which contributes to mitigating the effects of oxidative stress by activating the antioxidant system (Kour et al., 2023). After 5 days of water deprivation, we noted an increase in the levels of Na, K, and Ca, with a simultaneous decrease in the amount of Si, compared to the leaves of watered plants. After 20 days of cultivation in the water deficit conditions, the leaves were still characterised by increased content of only Na. In turn, an increase in the amount of K and Ca and a decrease in the level of Si were also noted in the root after 5 days of drought. However, in comparison to the leaves, Na remained at the same level. After 20 days, the levels of Na and K in the root also increased, compared to the control plants. It is undoubtedly possible to conclude that the disruption of the typical elemental composition is a result of drought. These findings highlight the plant’s adaptive response to drought, with alterations in its elemental composition that are a part of its strategy to cope with the stress. The increase in Na reflects the plant’s attempt to balance osmotic pressure under drought, where plant organ accumulates Na to maintain water balance and cell turgor in drought conditions. The rise in the Ca levels in the root indicates a response intended to maintain cellular function and stability of the egg-box in the cell wall under stress. In turn, the marked decrease in Si is related to structural changes in the cell wall (Saja-Garbarz et al., 2024) because Si accumulation in plants involves covalent bonding with cell wall components, i.e. pectin, potentially influencing wall structure and remodelling and thus creating a mechanical barrier (Coskun et al., 2019; Sheng and Chen, 2020). Si deficiency may potentially cause the observed changes in plant growth and physiology through lack of plasma membrane integrity, osmotic and salt stress tolerance, and nutrient homeostasis.

We can propose a conclusion that, upon alteration, the elemental signalling process is activated, which triggers subsequent adaptive responses, i.e., de novo synthesis or modification of the molecular properties of cell components, leading to cell wall remodelling. We obtained some clues to understanding structural alterations in plants. Firstly, this study revealed changes in the presence of unesterified homogalacturonans in cell walls under drought stress. As is well known, the middle lamella constitutes a reservoir of water, preventing water loss in drought conditions, and is primarily filled with an amorphous gel, mainly consisting of pectins. A stress-induced higher level of unesterified homogalacturonans permits calcium cross-linking, which enhances cell wall rigidity and helps in intracellular water preservation. Additionally, the effect is intensified by structural disorders of the root that are related to the uptake and subsequent transport of elements. This disorder makes it impossible to create specific interactions in the cell wall. We conclude that water loss can be linked to changes in the organisation of pectins and their ability to interlink with ions, suggesting that wall stiffening is connected with drought tolerance and turgor maintenance.

## CONCLUSION

Molecular studies using organ-specific antibodies open the way to the analysis of drought response networks, which can be targeted by different approaches. Molecules such as AGP, HG, or arabinoxylans prove useful in fine-tuning drought response pathways. Also, the side-chain modifications identified in the current work represent promising targets for future research, as they potentially play pivotal roles in the restructuring of cell walls in drought conditions. The next step of the research will be to confirm the obtained results in *in planta* studies aimed at localising cellular compartments that are a pioneering component of the adaptive response to environmental stress. We suspect that the middle lamella has an extremely important function in maintaining the proper structure of the tissue; hence, a thorough *in situ* analysis is necessary. In turn, basic research on the physiology of plant drought response in laboratory experiments and the transfer of these findings to application studies, searching for preventive solutions, will facilitate the development of new strategies to counteract water shortages.

## STATEMENTS & DECLARATIONS

### AUTHOR CONTRIBUTIONS

A.L. Conceptualisation, Formal analysis, Investigation, Methodology, Visualization, Writing – original draft; N.K. Investigation, Methodology, Data curation, Visualisation, Writing – review and editing

### CONFLICT OF INTEREST STATEMENT

No conflict of interest declared.

### FUNDING STATEMENT

This research received no specific grant from any funding agency in the public, commercial or not-for-profit sectors.

### DATA AVAILABILITY STATEMENT

Data will be shared on request to the corresponding author.

## Notes

### Competing Interest Statement

The authors have declared no competing interest.

## REFERENCES

Bowne JB, Erwin TA, Juttner J, et al. (2012). Drought responses of leaf tissues from wheat cultivars of differing drought tolerance at the metabolite level. Mol Plant 5: 418–429.

Bradford MM. (1976). A rapid and sensitive method for the quantitation of microgram quantities of protein utilizing the principle of protein-dye binding. Anal Biochem 72: 248–254.

Calderone S, Mauri N, Manga-Robles A, et al. (2024). Diverging cell wall strategies for drought adaptation in two maize inbreds with contrasting lodging resistance. Plant Cell Environ 47: 1747–1768.

Cantó-Pastor A, Kajala K, Shaar-Moshe L, et al. (2024). A suberized exodermis is required for tomato drought tolerance. Nat Plants 10: 118–130.

Chen P, Jung NU, Giarola V, et al. (2020). The dynamic responses of cell walls in resurrection plants during dehydration and rehydration. Front Plant Sci 10: 1698.

Cook BI, Mankin JS, Anchukaitis KJ. (2018). Climate change and drought: From past to future. Curr Clim Change Rep 4: 164–179.

Coskun D, Deshmukh R, Sonah H, et al. (2019). The controversies of silicon’s role in plant biology. New Phytol 221: 67–85.

Davies HA, Daniels MJ, Dow JM. (1997). Induction of extracellular matrix glycoproteins in Brassica petioles by wounding and in response to Xanthomonas campestris. MPMI 10: 812–820.

Ezquer I, Salameh I, Colombo L, et al. (2020). Plant cell walls tackling climate change: Insights into plant cell wall remodeling, its regulation, and biotechnological strategies to improve crop adaptations and photosynthesis in response to global warming. Plants 9: 212.

Fleck AT, Nye T, Repenning C, et al. (2011). Silicon enhances suberization and lignification in roots of rice (Oryza sativa). J Exp Bot 62: 2001–2011.

Ganie SA, Ahammed GJ. (2021). Dynamics of cell wall structure and related genomic resources for drought tolerance in rice. Plant Cell Rep 40: 437–459.

Gupta A, Rico-Medina A, Caño-Delgado AI. (2020). The physiology of plant responses to drought. Science 368: 266–269.

Hasanuzzaman M, Bhuyan MHMB, Nahar K, et al. (2018). Potassium: A vital regulator of plant responses and tolerance to abiotic stresses. Agron 8, 31.

Kim JS, Kidokoro S, Yamaguchi-Shinozaki K, et al. (2024). Regulatory networks in plant responses to drought and cold stress. Plant Physiol 195: 170–189.

Knox JP, Linstead PJ, Peart J, et al. (1991). Developmentally regulated epitopes of cell surface arabinogalactan proteins and their relation to root tissue pattern formation. Plant J 1: 317–326.

Kour J, Khanna K, Singh AD, et al. (2023). Calcium’s multifaceted functions: From nutrient to secondary messenger during stress. S Afr J Bot 152: 247–263.

Kutyrieva-Nowak N, Leszczuk A, Zdunek A. (2023). A practical guide to in situ and ex situ characterisation of arabinogalactan proteins (AGPs) in fruits. Plant Met 19: 117.

Li Z, Peng Y, Huang B. (2020). Transcriptional regulation of hydrogen peroxide and calcium for signaling transduction and stress-defensive genes contributing to improved drought tolerance in creeping bentgrass. J Am Soc Hortic Sci 145: 236–246.

Li WQ, Zhang MJ, Gan PF, et al. (2017). CLD1/SRL1 modulates leaf rolling by affecting cell wall formation, epidermis integrity and water homeostasis in rice. Plant J 92: 904–923.

Liu H, Ma Y, Chen N, et al. (2014). Overexpression of stress-inducible OsBURP16, the β subunit of polygalacturonase 1, decreases pectin content and cell adhesion and increases abiotic stress sensitivity in rice. Plant Cell Environ 37: 1144–1158.

Luquet D, Clément-Vidal A, Fabre D, et al. (2008). Orchestration of transpiration, growth and carbohydrate dynamics in rice during a dry-down cycle. Funct Plant Biol 35: 689–704.

Malik MA, Wani AH, Mir SH, et al. (2021). Elucidating the role of silicon in drought stress tolerance in plants. Plant Physiol Biochem 165: 187–195.

Marcus SE, Blake AW, Benians TAS, et al. (2010). Restricted access of proteins to mannan polysaccharides in intact plant cell walls. Plant J 64: 191–203.

McCartney L, Marcus SE, Knox JP. (2005). Monoclonal antibodies to plant cell wall xylans and arabinoxylans. J Histochem Cytochem 53: 543–546.

Moller I, Marcus SE, Haeger A, et al. (2008). High-throughput screening of monoclonal antibodies against plant cell wall glycans by hierarchical clustering of their carbohydrate microarray binding profiles. Glycoconjugate J 25: 37–48.

Moore JP, Vicré-Gibouin M, Farrant JM, et al. (2008). Adaptations of higher plant cell walls to water loss: drought vs desiccation. Physiol Plant 134: 237–245.

Naumann G, Cammalleri C, Mentaschi L, et al. (2021). Increased economic drought impacts in Europe with anthropogenic warming. Nat Clim Change 11: 485–491.

Oddo E, Inzerillo S, La Bella F, et al. (2011). Short-term effects of potassium fertilization on the hydraulic conductance of Laurus nobilis L. Tree Physiol 31: 131–138.

Oomen RJFJ, Doeswijk-Voragen CHL, Bush MS, et al. (2002). In muro fragmentation of the rhamnogalacturonan I backbone in potato (Solanum tuberosum L.) results in a reduction and altered location of the galactan and arabinan side-chains and abnormal periderm development. Plant J 30: 403–413.

Rakszegi M, Lovegrove A, Balla K, et al. (2014). Effect of heat and drought stress on the structure and composition of arabinoxylan and β-glucan in wheat grain. Carbohydr Polym 102: 557–565.

Saja-Garbarz D, Godel-Jędrychowska K, Kurczyńska E, et al. (2024). The effect of silicon supplementation and drought stress on the deposition of callose and chemical components in the cell walls of the Brassica napus roots. BMC Plant Biol 24: 1249.

Sheng H, Chen S. (2020). Plant silicon-cell wall complexes: Identification, model of covalent bond formation and biofunction. Plant Physiol Biochem 155: 13–19.

Skirycz A, Inzé D. (2010). More from less: plant growth under limited water. Curr Opin Biotechnol 21: 197–203.

Smallwood M, Martin H, Knox JP. (1995). An epitope of rice threonine- and hydroxyproline-rich glycoprotein is common to cell wall and hydrophobic plasma-membrane glycoproteins. Planta 196: 510–522.

Song WY, Zhang ZB, Shao HB, et al. (2008). Relationship between calcium decoding elements and plant abiotic-stress resistance. Int J Biol Sci 4: 116–125.

Stitt M, Gibon Y, Lunn JE, et al. (2007). Multilevel genomics analysis of carbon signalling during low carbon availability: coordinating the supply and utilisation of carbon in a fluctuating environment. Funct. Plant Biol 34: 526–549.

Tenhaken R. (2015). Cell wall remodeling under abiotic stress. Front. Plant Sci 5: 771.

Torun H. (2022). Combined efficiency of salicylic acid and calcium on the antioxidative defense system in two different carbon-fixative turfgrasses under combined drought and salinity. S Afr J Bot 144: 72–82.

Vaahtera L, Schulz J, Hamann T. (2019). Cell wall integrity maintenance during plant development and interaction with the environment. Nat Plants 5: 924–932.

Valarmathi R, Mahadeva Swamy HK, Appunu C, et al. (2023). Comparative transcriptome profiling to unravel the key molecular signalling pathways and drought adaptive plasticity in shoot borne root system of sugarcane. Sci Rep 13: 12853.

Verelst W, Bertolini E, De Bodt S, et al. (2013). Molecular and physiological analysis of growth-limiting drought stress in Brachypodium distachyon leaves. Mol Plant 6: 311–322.

Verhertbruggen Y, Marcus SE, Haeger A, et al. (2009a). An extended set of monoclonal antibodies to pectic homogalacturonan. Carbohydr. Res 344: 1858–1862.

Verhertbruggen Y, Marcus SE, Haeger A, et al. (2009b). Developmental complexity of arabinan polysaccharides and their processing in plant cell walls. Plant J 59: 413–425.

de Vries FT, Griffiths RI, Knight CG, et al. (2020). Harnessing rhizosphere microbiomes for drought-resilient crop production. Science 368: 270–274.

Wang X, Li X, Zhao W, et al. (2024). Current views of drought research: experimental methods, adaptation mechanisms and regulatory strategies. Front Plant Sci 15: 1371895.

Wang M, Wang R, Mur LAJ, et al. (2021). Functions of silicon in plant drought stress responses. Horti Res 8: 254.

Willats WGT, Orfila C, Limberg G, et al. (2001). Modulation of the degree and pattern of methyl-esterification of pectic homogalacturonan in plant cell walls: Implications for pectin methyl esterase action, matrix properties, and cell adhesion. JBC 276: 19404–19413.

Yang H, Nukunya K, Ding Q, et al. (2022). Tissue-specific transcriptomics reveal functional differences in floral development. Plant Physiol 188: 1158–1173.

Yang L, Wang CC, Guo WD, et al. (2006). Differential expression of cell wall related genes in the elongation zone of rice roots under water deficit. Russ J Plant Physiol 53: 390–395.

Yates EA, Valdor J-F, Haslam SM, et al. (1996). Characterization of carbohydrate structural features recognized by anti-arabinogalactan-protein monoclonal antibodies. Glycobiol 6: 131–139.

Zhang D, Tian C, Mai W. (2024). Exogenous sodium and calcium alleviate drought stress by promoting the succulence of Suaeda salsa. Plants 13: 721.

Zhao N, Zhou Z, Cui S, et al. (2024). Selected cell wall remodeling mechanisms orchestrating plant drought tolerance. Plant Stress, 100698.

Zhu Y, Guo J, Feng R, et al. (2016). The regulatory role of silicon on carbohydrate metabolism in Cucumis sativus L. under salt stress. Plant Soil 406: 231–249.

